# Mycobacterial Serine/Threonine phosphatase PstP is phospho-regulated and localized to mediate control of cell wall metabolism

**DOI:** 10.1101/2022.02.28.482390

**Authors:** Farah Shamma, E. Hesper Rego, Cara C. Boutte

## Abstract

The mycobacterial cell wall is profoundly regulated in response to environmental stresses, and this regulation contributes to antibiotic tolerance. The reversible phosphorylation of different cell wall regulatory proteins is a major mechanism of cell wall regulation. Eleven Serine/Threonine protein kinases (STPKs) phosphorylate many critical cell wall-related proteins in mycobacteria. PstP is the sole serine/ threonine phosphatase, but few proteins have been verified as PstP substrates. PstP is itself phosphorylated but the role of its phosphorylation in regulating its activity has been unclear. In this study we aim to discover novel substrates of PstP in *Mycobacterium tuberculosis* (*Mtb*). We show *in vitro* that PstP dephosphorylates two regulators of peptidoglycan in *Mtb*, FhaA and Wag31. We also show that a phospho-mimetic mutation of T137 on PstP negatively regulates its catalytic activity against the cell wall regulators FhaA, Wag31, CwlM, PknB and PknA, and that the corresponding mutation in *Mycobacterium smegmatis (Msmeg)* causes mis-regulation of peptidoglycan *in vivo*. We show that PstP is localized to the septum, which likely restricts its access to certain substrates. These findings on the regulation of PstP provide insight into the control of cell wall metabolism in mycobacteria.

## INTRODUCTION

*Mycobacterium tuberculosis* (*Mtb*), which causes Tuberculosis (TB), remains a public health menace, infecting about 10 million and killing at least a million people worldwide each year (World Health Organization, 2021). TB infections require extended multi-drug antibiotic treatment regimens (Nguyen, 2016). The mycobacterial cell wall contributes to *Mtb*’s inherent antibiotic tolerance (Jarlier and Nikaido, 1994; Sarathy *et al*., 2013; Batt *et al*., 2020). The wall serves as permeability barrier (Jarlier and Nikaido, 1990; Hett and Rubin, 2008; Hoffmann *et al*., 2008; Sarathy *et al*., 2013), and is strongly regulated in response to stresses (Cunningham and Spreadbury, 1998; Betts *et al*., 2002; Bhamidi *et al*., 2012; Sarathy *et al*., 2013), including infection (Sharma *et al*., 2006). The cell wall changes in stress are associated with increased antibiotic tolerance (Xie *et al*., 2005; Liu *et al*., 2016; Sarathy *et al*., 2017). The expression and activity of many cell wall enzymes and regulators are also tightly coordinated over the course of the cell cycle in order to maintain cell wall integrity during elongation and division (Kieser and Rubin, 2014; Dulberger *et al*., 2020; Bandekar *et al*., 2020).

The cell wall consists of three covalently linked layers: a peptidoglycan layer surrounding the plasma membrane, an arabinogalactan layer and a mycolic acid layer (Marrakchi *et al*., 2014). Peptidoglycan is a network of glycan chains cross-linked by small peptides, and is the innermost layer of the wall (Abrahams and Besra, 2021). Reversible Serine/Threonine (S/T) phosphorylation of cell wall enzymes and regulators is critical for regulation of cell wall metabolism in response to environmental signals (Wang *et al*., 1998; Juris *et al*., 2000; Echenique *et al*., 2004). There are 11 Serine/Threonine kinases (STPKs) (PknA-B and PknD-L) and one essential S/T protein phosphatase PstP in *Mtb* (Kang *et al*., 2005; Gee *et al*., 2012; Baer *et al*., 2014; Boutte *et al*., 2016)(Cole *et al*., 1998; Bach *et al*., 2009). PstP is also essential in *Msmeg* (Sharma *et al*., 2016; DeJesus *et al*., 2017).

PstP belongs to the protein phosphatase 2C (PP2C) subfamily of divalent metal-ion dependent protein serine/threonine phosphatases (Cohen, 1989; Barford, 1996; Chopra *et al*., 2003), which regulate prokaryotic and eukaryotic cell growth and division, sporulation, stress response and metabolic processes in response to various environmental signals (Vijay *et al*., 2000:2; Irmler and Forchhammer, 2001; Mougous *et al*., 2007; Lu and Wang, 2008; Bradshaw and Losick, 2015; Bradshaw *et al*., 2017). PstP*_Mtb_* shares structural architecture and conserved catalytic residues with the human PP2Cα (Pullen *et al*., 2004) (Figure 1), the PP2C family prototype. The N-terminal cytoplasmic enzymatic domain of PstP*_Mtb_* (Figure 1) connects to its C-terminal extracellular domain via a transmembrane helix (Pullen *et al*., 2004). PstP*_Mtb_* is itself phosphorylated on T137, T141, T174 and T290 (Figure 1 (Sajid *et al*., 2011). Phosphorylation on all these sites together affects PstP activity against small molecule substrates *in vitro* (Sajid *et al*., 2011). And phosphorylation on T174 affects cell wall metabolism *in vivo* in *Msmeg* (Shamma *et al*., 2021).

**Figure 1.**
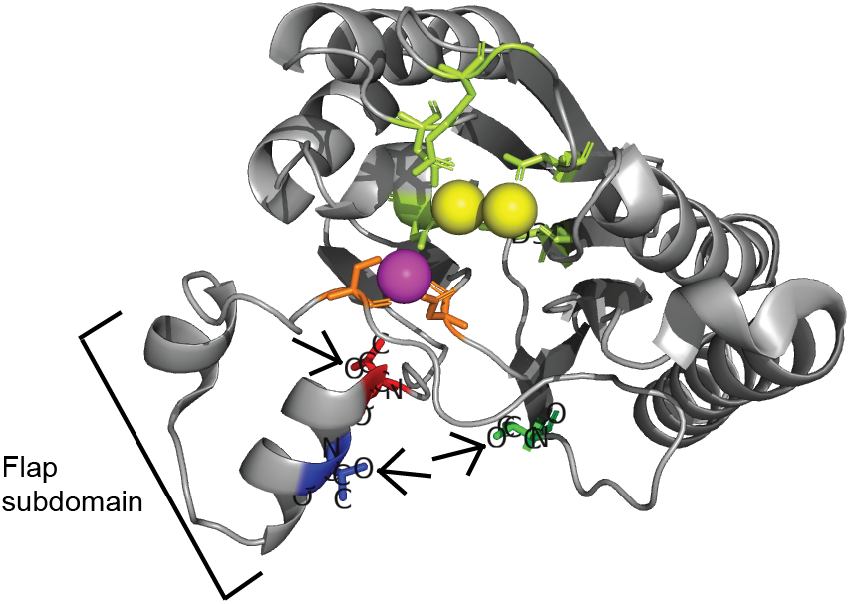
Phosphorylation sites are not near the active site in PstP. Schematic of the crystal structure of PstP from *M. tuberculosis* (PstPWT*_Mtb_*) (Pullen *et al*., 2004) (PDB code 1TXO) showing the conserved threonines (T) which are phosphorylated by the kinases PknA and PknB (Sajid *et al*., 2011). Highlighted on the structure are: red -T137 (T134 in PstP*_Msmeg_*); blue - T141 (T138 in PstP*_Msmeg_*); green - T174 (T171 in PstP*_Msmeg_*); yellow - Mn^2+^ present at the conserved two-metal center typically seen in the PP2C subfamily of protein Serine/Threonine phosphatases, which are part of the active site core; pink - the unique third Mn^2+^ ion bound in the catalytic core of PstP*_Mtb_* but not present in the human PP2Cα which sits in a groove created by the flap subdomain next to the active site; orange - Ser60 and Asp118 that act as ligands for the third Mn ion; lime green - residues of the catalytic sites surrounding the two-metal center. Arrows point to position of the OH-groups on T137, T141 and T174 that are phosphorylated.

As PstP is the only mycobacterial S/T protein phosphatase (Cole *et al*., 1998), it likely dephosphorylates many of the substrates phosphorylated by the 11 STPKs (Prisic and Husson, 2014). However, only a few proteins are known PstP substrates: the STPKs PknA and PknB (Boitel *et al*., 2003; Chopra *et al*., 2003; Durán *et al*., 2005; Sajid *et al*., 2011), mycolic acid biosynthesis enzymes KasA and KasB (Molle *et al*., 2006), the arabinogalactan regulator EmbR (Sharma *et al*., 2006), and CwlM (Shamma *et al*., 2021), which in its phosphorylated form is an essential activator of MurA (Boutte *et al*., 2016), the first enzyme of peptidoglycan precursor synthesis (Marquardt *et al*., 1992). So dephosphorylation of CwlM∼P by PstP (Shamma *et al*., 2021) should halt peptidoglycan synthesis. CwlM∼P also interacts with FhaA (Turapov *et al*., 2018), which is involved in modulating peptidoglycan biosynthesis (Gee *et al*., 2012; Viswanathan *et al*., 2015; Viswanathan *et al*., 2017) and unphosphorylated CwlM interacts with the peptidoglycan precursor lipid-II flippase MurJ (Turapov *et al*., 2018), suggesting that CwlM may regulate multiple steps in peptidoglycan synthesis (Turapov *et al*., 2018).

FhaA and Wag31 are both regulators of mycobacterial cell wall metabolism. FhaA helps maintain cell envelope integrity, cell length and antibiotic tolerance (Viswanathan *et al*., 2015; Viswanathan *et al*., 2017). FhaA stabilizes PbpA (Viswanathan *et al*., 2017) and binds and may inhibit the MurJ flippase (Gee *et al*., 2012). FhaA is phosphorylated by PknB (Grundner *et al*., 2005; Prisic *et al*., 2010; Roumestand *et al*., 2011), though the role of its phosphorylation is not yet clear. Wag31 localizes at the cell poles (Kang *et al*., 2005), is essential for polar peptidoglycan synthesis (Kang *et al*., 2008; Jani *et al*., 2010) and is phosphorylated by the STPK PknA (Kang *et al*., 2005). Wag31’s phosphorylation may also have a role in controlling peptidoglycan metabolism (Jani *et al*., 2010). It was not known whether FhaA and Wag31 were actively dephosphorylated, or if their phosphorylated forms were removed through protein degradation.

In this study we show that FhaA*_Mtb_* and Wag31*_Mtb_* are substrates of PstP*_Mtb_ in vitro*, and that PstP dephosphorylates Wag31 much faster than it does FhaA or CwlM. We show that the T137E phospho-mimetic mutation of PstP*_Mtb_* inhibits PstP*_Mtb_*’s catalytic activity against the substrates CwlM, FhaA, Wag31, PknB and PknA *in vitro*. Furthermore, the corresponding phospho-mimetic mutation in *Msmeg* leads to misregulation of peptidoglycan metabolism. We also find that PstP is enriched at the cell division site, suggesting that its access to substrates might also control its activity. We suggest a model where PstP’s activity against assorted substrates is regulated by substrate affinity and protein localization, and further toggled by the phosphorylations on PstP.

## RESULTS

### PstP*_Mtb_* dephosphorylates FhaA*_Mtb_* and Wag31*_Mtb_ in vitro*

FhaA*_Mtb_* is phosphorylated by PknB*_Mtb_ in vivo* (Prisic *et al*., 2010) and *in vitro* (Grundner *et al*., 2005; Roumestand *et al*., 2011) on T116 (Roumestand *et al*., 2011). Wag31*_Mtb_* is phosphorylated at T73 by PknA*_Mtb_* (Kang *et al*., 2005). To determine if FhaA*_Mtb_* and Wag31*_Mtb_* are substrates of PstP*_Mtb_*, we purified full-length His-FhaA*_Mtb_*, His-Wag31*_Mtb_* and phosphorylated them by by His-MBP fusions to the kinase domains of PknA*_Mtb_* and PknB*_Mtb_*. We then added the cytosolic domain of PstP*_Mtb_* containing the catalytic domain (His-PstP_c_WT*_Mtb_*) and measured changes in phosphorylation. All the proteins in this study were purified first by nickel column and then by size exclusion chromatography, to ensure proper folding. In all the kinase reaction mixtures in this study, the molar concentrations of the substrates were in excess over the ATP [γ-^32^P], so that there is no leftover ATP [γ-^32^P] to allow re-phosphorylation once His-PstP_c_ is added to the reaction mixture for the phosphatase assay. This experimental setup allows us to measure dephosphorylation in the presence of the kinases without re-phosphorylation happening simultaneously.

We phosphorylated His-FhaA*_Mtb_* with His-MBP-PknB*_Mtb_* and ATP [γ-^32^P]. After addition of His-PstP_c_WT*_Mtb_*, at 1/10^th^ the concentration of the His-FhaA*_Mtb,_* the phosphorylation on His-FhaA*_Mtb_*∼P decreased by ∼80% in 80 minutes (Figure 2a,b). Our control assay with no phosphatase shows that the phosphorylation on His-FhaA*_Mtb_*∼P is stable over time in the phosphatase buffer (Figure 2a, b). We then phosphorylated His-Wag31*_Mtb_* with His-MBP-PknA*_Mtb_* and ATP [γ-^32^P], and added His-PstP_c_WT*_Mtb_*. With 1/100^th^ the amount of PstP compared to Wag31, Wag31*_Mtb_* phosphorylation decreased by about ∼90% in 2 minutes (Figure 3a,b), only in the presence of His-PstP_c_WT*_Mtb_*. These data show that FhaA*_Mtb_* and Wag31*_Mtb_* are substrates of PstPWT*_Mtb_ in vitro*.

**FIGURE 2.**
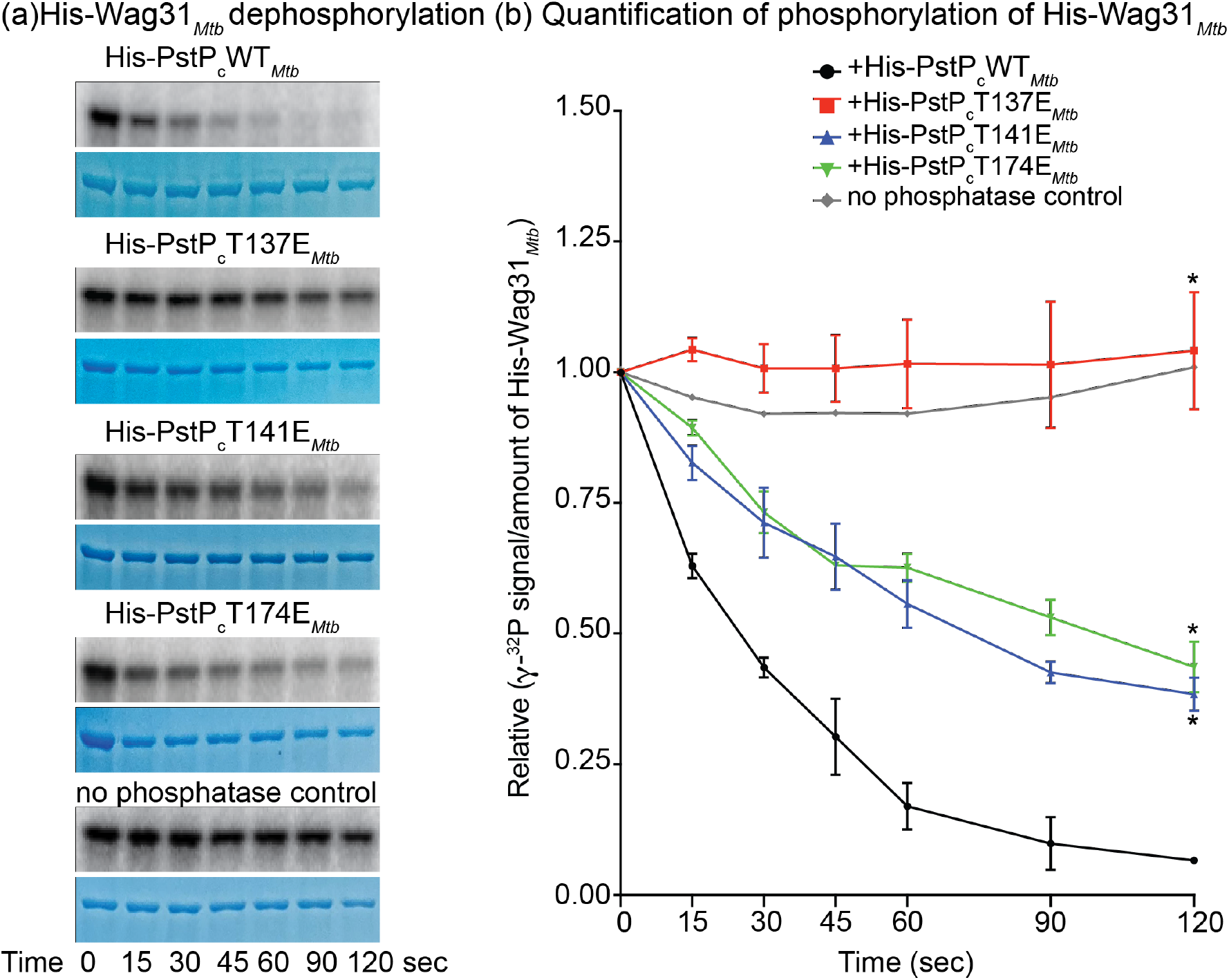
Dephosphorylation of FhaA by PstP and phospho-mimetic isoforms. (a) Autoradiograms and Coomassie-stained SDS gels of *in vitro* phosphatase assays performed with ATP-[γ-^32^P]-phosphorylated His-FhaA*_Mtb_* and His-PstP_c_WT*_Mtb_* (first panel), His-PstP_c_137E*_Mtb_* (second panel), His-PstP_c_T141E*_Mtb_* (third panel), His-PstP_c_T174E*_Mtb_* (fourth panel) and no phosphatase control reaction (bottom panel). The ratio of each phosphatase to His-FhaA*_Mtb_* in each of the assays was 1:10. One set of representative images for each assay is shown here from two individual assays performed with individually purified batches of phosphatases. (b) Quantification of relative intensities of [γ-^32^P] on His-FhaA*_Mtb_* on autoradiogram over the relative amount of His-FhaA*_Mtb_* on the SDS gel, average between two replicate experiments. The relative values refer to the normalized values of intensity of [γ-^32^P] signal and the protein bands on gel at each time point (0 to 80min) with the 0 time point. P values were calculated using two-tailed unpaired t-test. P-values of T137E at 0 min *vs.* T137E 80 min =0.7598, WT *vs.* T137E at 80 min= 0.0487, WT *vs.* T141E at 80 min =0.3712, WT *vs.* T174E at 80 min=0.2000. The error bars represent standard error of mean.

**FIGURE 3.**
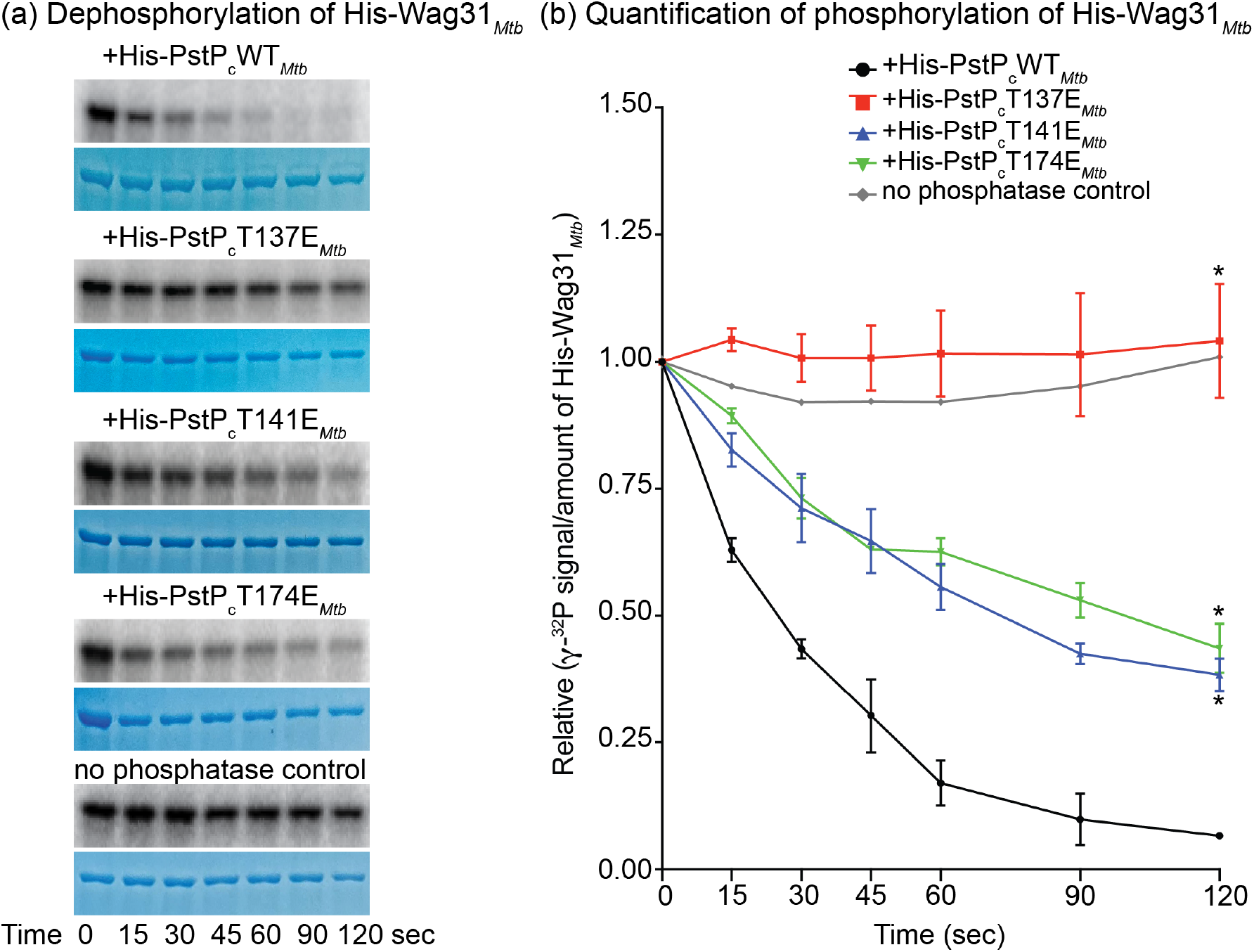
Dephosphorylation of Wag31*_Mtb_* by PstP and phospho-mimetic isoforms. (a) Autoradiograms and Coomassie-stained SDS gels of *in vitro* phosphatase assays performed with ATP-[γ-^32^P]-phosphorylated His-Wag31*_Mtb_* and His-PstP_c_WT*_Mtb_* (first panel), His-PstP_c_WT*_Mtb_* (second panel), His-PstP_c_T141E*_Mtb_* (third panel), His-PstP_c_T174E*_Mtb_* (fourth panel) and no phosphatase control reaction (bottom panel). The ratio of each phosphatase to His-Wag31*_Mtb_* in the assays was 1:100. One set of representative images for each reaction is shown here from two individual assays performed with individually purified batches of phosphatases. (b) Quantification of relative intensities of [γ-^32^P] on His-Wag31*_Mtb_* on autoradiograms over the relative amount of His-Wag31*_Mtb_* on the SDS gel, average between two replicate experiments. The relative values are the normalized values of intensity of [γ-^32^P] signal and the protein bands on gel at each time point (0 to 120 sec) with the 0 time point. P values were calculated using two-tailed unpaired t-test. P-values of T137E at 0 sec *vs.* 15 sec = 0. 0.1947 and 0 sec *vs.* 120sec= 0.7493, WT *vs.* T137E at 120 sec= 0.0130, WT *vs.* T141E at 120 sec =0.0106 and WT *vs.* T174E at 120 sec = 0.170. The error bars represent standard error of mean.

Our previous data that demonstrated PstP*_Mtb_* dephosphorylates CwlM*_Mtb_* (Shamma *et al*., 2021) is reproducible in our *in vitro* assays with ATP [γ-^32^P] where we observe a 97% decrease in phosphorylation on His-SUMO-CwlM*_Mtb_*∼P 40 minutes after addition of 1/10^th^ the amount of His-PstP_c_WT*_Mtb_* (Figure 4a, b). In all the assays, we see no phosphorylation signals on the WT and phospho-mimetic PstPs on the autoradiograms (Supplementary figure S3). This is likely because PstP can quickly dephosphorylate itself (Sajid *et al*., 2011) and the purified forms are thus unphosphorylated.

**FIGURE 4.**
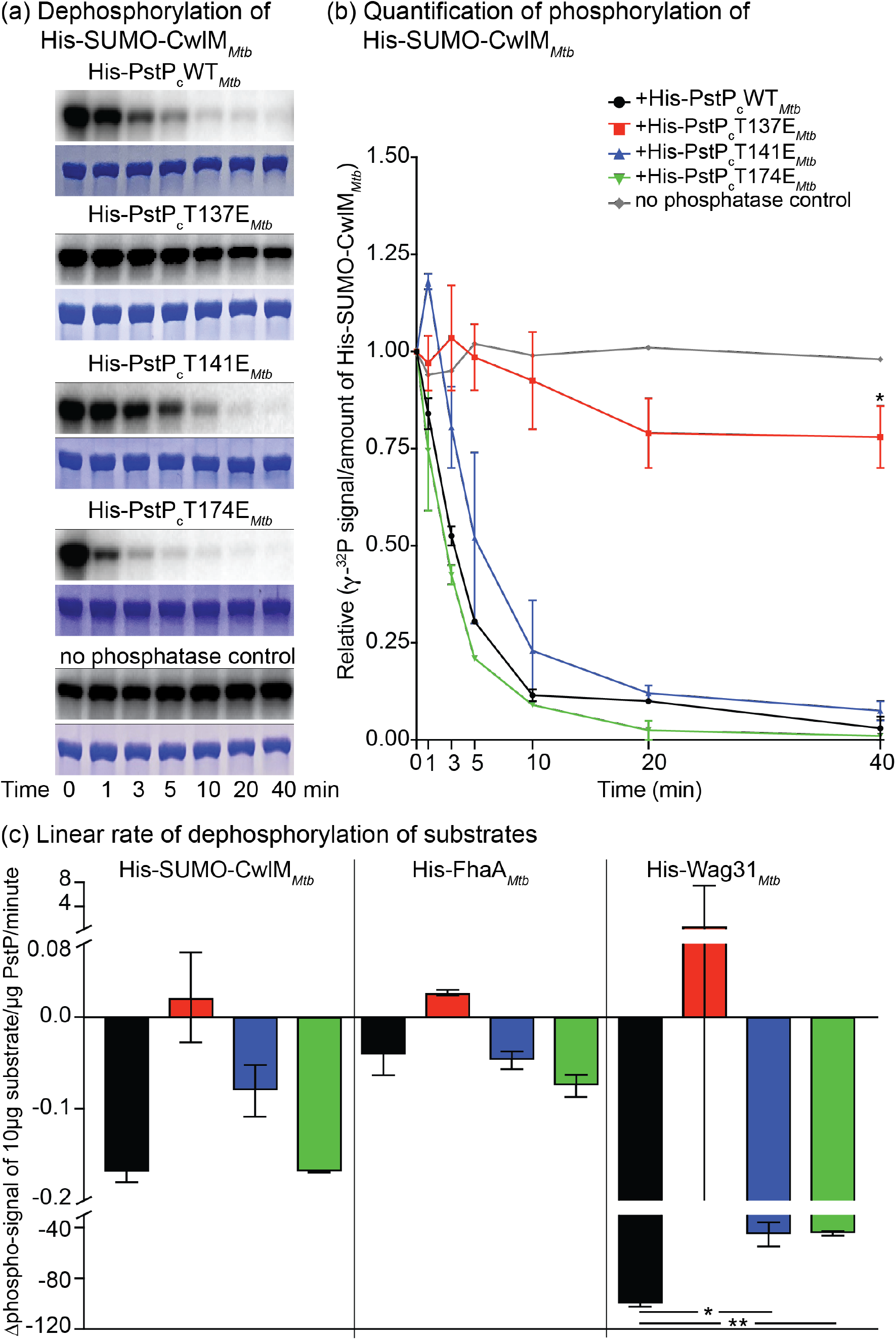
Dephosphorylation of CwlM*_Mtb_* by PstP*_Mtb_* its phospho-mimetic isoforms and linear rate of dephosphorylation of CwlM*_Mtb_*, FhaA*_Mtb_* and Wag31*_Mtb_*. (a) Autoradiograms and Coomassie-stained SDS gels of *in vitro* phosphatase assays performed with ATP-[γ-^32^P]-phosphorylated His-SUMO-CwlM*_Mtb_* and His-PstP_c_WT*_Mtb_* (first panel), His-PstP_c_WT*_Mtb_* (second panel), His-PstP_c_T141E*_Mtb_* (third panel), His-PstP_c_T174E*_Mtb_* (fourth panel) and no phosphatase control reaction (bottom panel). The ratio of each phosphatase to His-SUMO-CwlM*_Mtb_* was 1:10. One set of representative images for each reaction is shown here from two individual assays performed with individually purified batches of phosphatases. (b) Quantification of relative intensities of [γ-^32^P] on His-SUMO-CwlM*_Mtb_* on autoradiogram over the relative amount of His-SUMO-CwlM*_Mtb_* on the SDS gel, average between two replicate experiments. The relative values are the normalized values of intensity of [γ-^32^P] signal and the protein bands on gel at each time point (0 to 40min) with the 0 time point. P values were calculated using two-tailed unpaired t-test. P-values of T137E at 0 min *vs.* 40 min= 0.1107, WT *vs*. T137E at 40 min= 0.0127, WT *vs*. T141E at 40 min=0.3683 and WT *vs*. T174E at 40 min= 0.5918. The error bars represent standard error of mean. (c) Quantification of linear rate of change in [γ-^32^P] signal on 10μg of His-SUMO-CwlM*_Mtb_*, His-FhaA*_Mtb_* and His-Wag31*_Mtb_* by 1μg His-PstP_c_WT*_Mtb_* and its phospho-mimetic mutants per minute. P values were calculated using two-tailed unpaired t-test. **, *P* value = 0.0026 and *, *P* value 0.0306. The error bars represent standard error of mean.

Because measuring the rates of dephosphorylation by PstP against different substrates required different PstP concentrations and timing, we calculated the initial linear rate of dephosphorylation per unit concentration of PstP, in order to compare PstP’s activity against different substrates. This comparison indicates that His-PstP_c_WT*_Mtb_* dephosphorylates His-Wag31*_Mtb_* more than 500 times faster than His-SUMO-CwlM*_Mtb_* and about 2500 times faster than His-FhaA*_Mtb_* (Figure 4c).

### Phospho-mimetic mutation at PstPT137*_Mtb_* inhibits phosphatase activity

PstP*_Mtb_* is phosphorylated on the conserved threonine (T) residues 137, 141 and 174 (Figure 1), and when phosphorylated, shows increased activity against small molecule substrates *in vitro* (Sajid *et al*., 2011). We hypothesized that phosphorylation of the threonine residues of PstP individually might help regulate PstP’s activity by either affecting its catalytic activity broadly, or by specifically affecting substrate affinity or activity against specific substrates. To test this, we made phospho-mimetic (T to E) (Cottin *et al*., 1999) mutants of PstP*_Mtb_* at each of the three conserved phosphorylation sites of PstP*_Mtb_* in the catalytic domain and purified them (His-PstP_c_T137E*_Mtb_*, His-PstP_c_T141E*_Mtb_* and His-PstP_c_T174E*_Mtb_*_)_. We previously showed that *in vitro* His-PstP_c_WT*_Mtb_* and His-PstP_c_T174E*_Mtb_* dephosphorylate His-SUMO-CwlM*_Mtb_* at the same rate (Shamma *et al*., 2021). We performed separate *in vitro* phosphatase assays with the substrates His-FhaA*_Mtb_*, His-Wag31*_Mtb_* and His-SUMO-CwlM*_Mtb_* with the phospho-mimetic versions of PstP. All three substrates exhibit stable phosphorylation with His-PstP_c_T137E*_Mtb_*, indicating that it is inactive *in vitro* against all three substrates (Figures 2,3,4). This suggests that phosphorylation on T137 of PstP may inhibit its activity against multiple substrates.

His-MBP-PknB*_Mtb_* and His-MBP-PknA*_Mtb_*, the STPKs used in these assays, are also confirmed substrates of PstP (Sajid *et al*., 2011). PknA and PknB are activated to phosphorylate their own substrates when they themselves are phosphorylated at their activation loops (Baer *et al*., 2014). Because we already had these kinases in our phosphatase assays, we were able to measure dephosphorylation of His-MBP-PknB*_Mtb_* and His-MBP-PknA*_Mtb_* by PstP in the context of a 10-fold and 5-fold excess of His-SUMO-CwlM*_Mtb_* or His-Wag31*_Mtb_*, respectively. We found that His-PstP_c_T137E*_Mtb_* dephosphorylates both His-MBP-PknB*_Mtb_* (Figure 5a,b) and His-MBP-PknA*_Mtb_* (Figure 5c,d) more slowly than His-PstP_c_WT*_Mtb_* and the other PstP mutants. This shows that His-PstP_c_T137E*_Mtb_* is inhibited against at least five different substrates.

**FIGURE 5.**
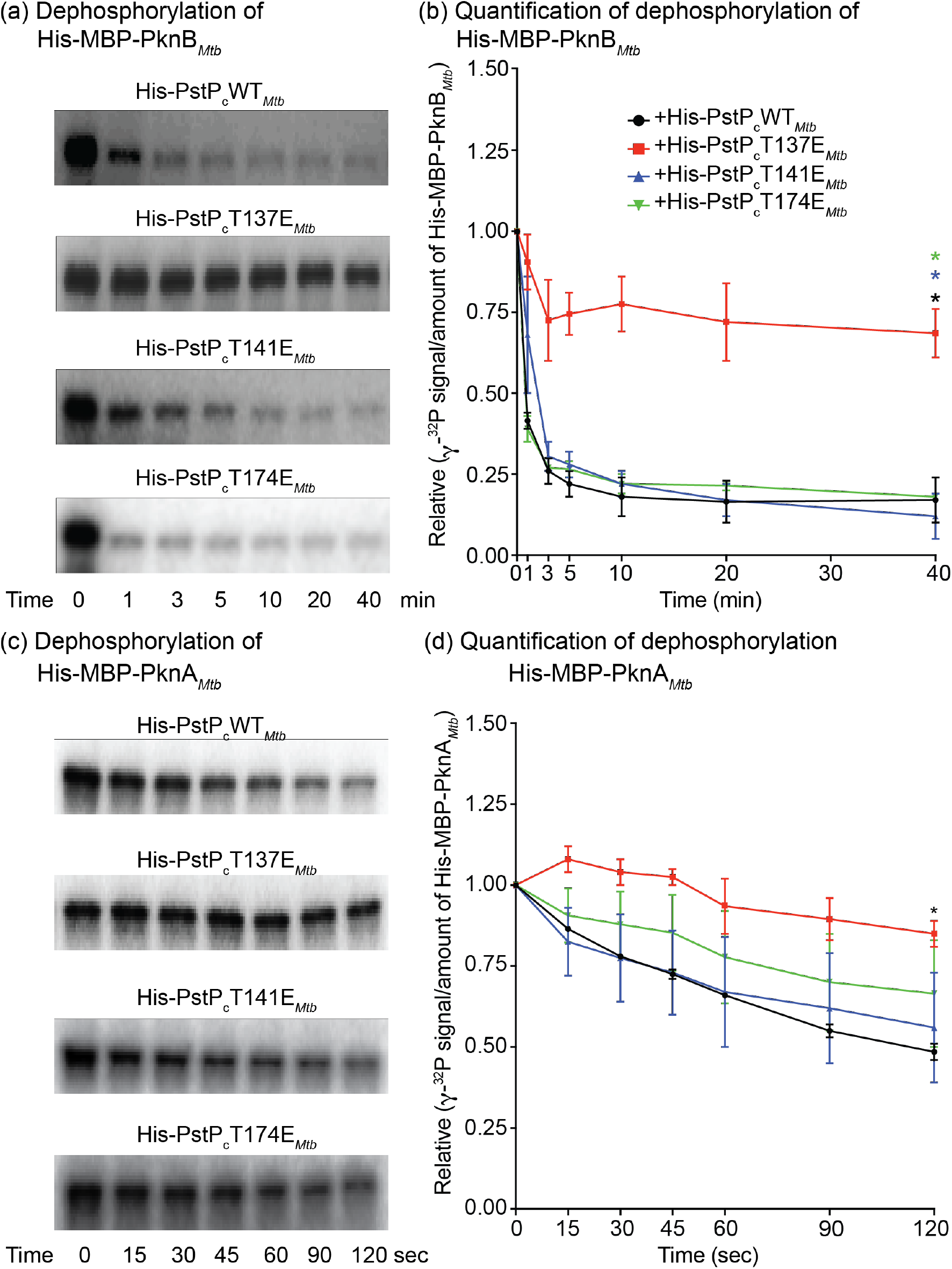
Dephosphorylation of PknB*_Mtb_* and PknA by PstPWT*_Mtb_* and its phospho-mimetic variants. (a) Autoradiograms showing [γ-^32^P]-phospho-signals on His-MBP-PknB*_Mtb_* in the *in vitro* phosphatase assays with His-SUMO-CwlM*_Mtb_* and His-PstP_c_WT*_Mtb_* (first panel), His-PstP_c_WT*_Mtb_* (second panel), His-PstP_c_T141E*_Mtb_* (third panel) and His-PstP_c_T174E*_Mtb_* (fourth panel), where His-MBP-PknB*_Mtb_* was used to phosphorylate His-SUMO-CwlM*_Mtb_*. One set of representative images for each reaction is shown here from two individual assays performed with individually purified batches of phosphatases. (b) Quantification of relative intensities of [γ-^32^P] on His-MBP-PknB*_Mtb_* on autoradiogram, average between two replicate experiments. The relative values refer to the normalized values of intensity of [γ-^32^P] signal at each time point with the 0 time point. P values were calculated using two-tailed unpaired t-test. P value of T137E at 0 min *vs.* 40 min= 0.0523. *, *P* value= 0.0375 (WT *vs*. T137E at 40 min); *, *P* value=0.0314 (T141E *vs*.T137E at 40 min); *, *P* value= 0.0217 (T174E *vs*.T137E at 40 min). The error bars represent standard error of mean. (c) Autoradiograms showing [γ-^32^P]-phospho-signals on His-MBP-PknA*_Mtb_* in the *in vitro* phosphatase assays with His-Wag31*_Mtb_* and His-PstP_c_WT*_Mtb_* (first panel), His-PstP_c_WT*_Mtb_* (second panel), His-PstP_c_T141E*_Mtb_* (third panel) and His-PstP_c_T174E*_Mtb_* (fourth panel), where His-Wag31*_Mtb_* was first phosphorylated with His-MBP-PknA*_Mtb_*. One set of representative images for each reaction is shown here from two individual assays performed with individually purified batches of phosphatases. (d) Quantification of relative intensities of [γ-^32^P] on His-MBP-PknA*_Mtb_* on autoradiogram. P values were calculated using two-tailed unpaired t-test. The relative values refer to the normalized values of intensity of [γ-^32^P] signal at each time point with the 0 time point. P value of T137E at 0 sec *vs.* 120 sec= 0.0643. *, *P* value= 0.0163 (WT *vs*. T137E at 120 sec). The error bars represent standard error of mean.

### Phospho-mimetic mutations at PstPT141*_Mtb_* and PstPT174*_Mtb_* moderately inhibit Wag31’s dephosphorylation

His-PstP_c_T141E*_Mtb_* and His-PstP_c_T174E*_Mtb_* dephosphorylate His-FhaA*_Mtb_* (Figure 2,4c) and His-SUMO-CwlM*_Mtb_* (Figure 4) at about the same rate as His-PstP_c_WT*_Mtb_*, but dephosphorylate His-Wag31*_Mtb_* about half as fast as His-PstP_c_WT*_Mtb_* (Figure 3,4c). These biochemical data support our hypothesis that phosphorylation on PstP may modulate its substrate specificity. Phosphorylation at T141 and T174 appear to inhibit binding or catalysis of Wag31 but not FhaA or CwlM.

### A negative charge on T134 of PstP*_Msmeg_* regulates peptidoglycan metabolism in starvation

Phosphorylation of CwlM allows it to activate the peptidoglycan precursor enzyme MurA, thereby licensing peptidoglycan synthesis (Boutte *et al*., 2016). Phosphorylation of CwlM also affects its binding to FhaA and the lipid II flippase MurJ (Turapov *et al*., 2018). CwlM is quickly dephosphorylated in the transition to stasis (Boutte *et al*., 2016) likely by PstP (Shamma *et al*., 2021). The function of phosphorylation on FhaA and Wag31 is not yet clear, but available data (Jani *et al*., 2010; Gee *et al*., 2012; Viswanathan *et al*., 2016) suggest it could also affect peptidoglycan metabolism. Dephosphorylation of these regulators could be important for downregulation of peptidoglycan metabolism in stasis.

As the phospho-mimetic mutation at PstPT137*_Mtb_* inhibits PstP’s dephosphorylation of three peptidoglycan regulatory proteins *in vitro* (Figure 2,3,4), we sought to determine if the corresponding phospho-site on PstP (T134) might affect peptidoglycan metabolism *in vivo* in *Msmeg*. Although this site is not known to be phosphorylated in PstP*_Msmeg_* (Iswahyudi *et al*., 2019; Le *et al*., 2020), the PstP proteins are 78% identical between *Msmeg* and *Mtb* and most of PstP’s substrates are also conserved, so we reasoned that performing experiments with a negative charge at residue T134 on PstP in *Msmeg* will provide a preliminary approximation of what the effect of phosphorylation on this residue might be in *Mtb.* We made *Msmeg* strains with a phospho-mimetic *pstP*T134E*_Msmeg_* allele and a phospho-ablative *pstP*T134A*_Msmeg_* allele replacing the wild-type *pstP*. We verified the expression of PstPT134E*_Msmeg_* and T134A in these strains under normal growth condition (Figure 6h). We found that PstPT134E*_Msmeg_* was stable but PstPT134A*_Msmeg_* was not, so we did not use the *pstP*T134A*_Msmeg_* allele strains for further experiments. We reported previously that the biological replicates of *pstP*T134E*_Msmeg_* mutant strains had inconsistent doubling times (Shamma *et al*., 2021) probably due to potential suppressor mutations. We confirmed that the slow growing clones could, after passaging, accumulate suppressor mutations that allowed them to grow as fast as the *pstP*WT*_Msmeg_* strain. To characterize the original *pstP*T134E*_Msmeg_* strain, we isolated numerous clones from fresh transformations, and measured the doubling times of each before continuing with experiments. We show data only from clones that grew at the slower rate.

**Figure 6.**
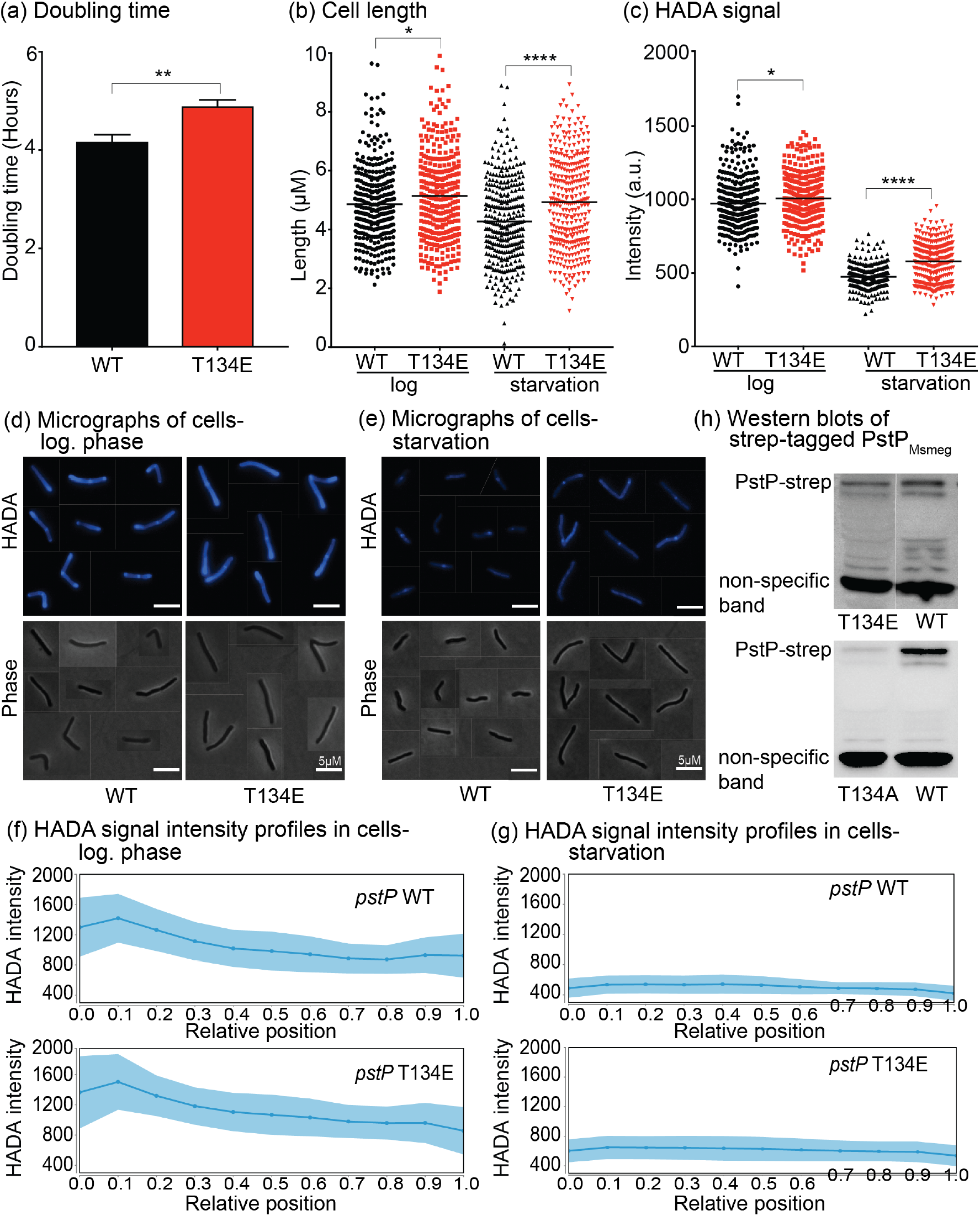
The phosphosite on T134 on PstP*_Msmeg_* affects PG metabolism. (a) Doubling times of *Msmeg* strains carrying the *pstP_Msmeg_* WT and the phospho-mimetic T134E (equivalent to T137 on PstP*_Mtb_*) alleles. Each bar represents the mean of individual average doubling times of three biological replicates of each genotype from two different experiments on different dates. The error bars represent mean with standard deviation. **, *P* value= 0.0026. (b) Quantification of cell lengths of isogenic *pstP_Msmeg_* allele strains (WT and T134E) grown in 7H9 in log. phase and starved in PBS with Tween80. At least 100 cells from each of three biological replicates were quantified. P values were calculated by unpaired t-test. *, *P* value = 0.0122, ****, *P* value= 0.000000031. (c) Quantification of mean intensities of HADA of *pstP_Msmeg_* allele strains (WT and T134E) grown in 7H9 in log. phase and starved in PBS with Tween80. Cells were stained after 15 minutes of starvation and washed and fixed after another 15 minutes (starved for a total of 30 minutes). At least 100 cells from each of three biological replicates were quantified. P values were calculated by unpaired t-test. *, *P* value = 0.0109, ****, *P* value= <0.000000000000001. (d) and (e) Representative HADA and phase images of cells from *pstP_Msmeg_* allele strains (WT and T134E) grown in 7H9 in log. phase (d) and after 30 minutes of starvation in PBS with Tween80 (e). (f) and (g) Intensity profiles of HADA signals in cells of *pstP_Msmeg_* allele strains (WT and T134E) in log. phase (f) and after starving for 30 minutes in PBS supplemented with Tween80 (g). The shaded regions denote standard deviations and the solid lines represent the mean intensity values. Signal intensities from at least 110 cells from each of three biological replicates of every *pstP_Msmeg_* allelic variant genotype were quantified and used to analyze the signal profiles in MicrobeJ. 0 to 1 on the X-axes represent cells from the old pole to the new pole, respectively. (h) α-strep western blots of strep-tagged PstP*_Msmeg_* in *pstP_Msmeg_*WT, *pstP_Msmeg_*T134E and *pstP_Msmeg_*T134A allele strains in log. phase. A non-specific band is shown as a loading control.

We stained both log. phase and starved *pstP*T134E*_Msmeg_* and WT*_Msmeg_* strains using a fluorescent D-amino acid HADA (Kuru *et al*., 2012) as a reporter of active peptidoglycan metabolism. We saw a slight increase in length (Figure 6b,d and Supplementary Figure S2a,b) and HADA intensity (Figure 6c,d and Supplementary Figure S2a,b) in the *pstP*T134E*_Msmeg_* strains in the log. phase compared to the WT*_Msmeg_* strains. After starvation, these differences increased (Figure 6b,c,e and Supplementary Figure S2c,d). WT*_Msmeg_* strains quickly downregulate peptidoglycan metabolism upon starvation (Boutte *et al*., 2016), as the decreased HADA staining (Figure 6c,d,e,f,g) shows. The starved *pstP*T134E*_Msmeg_* strains show significantly higher HADA signal than the WT*_Msmeg_* (Figure 6c, top right panels of d,e and Supplementary Figure S2b,d) indicating a defect in downregulating peptidoglycan metabolism. This suggests that the presence of a constant negative charge on phosphosite T134 of PstP*_Msmeg_* impairs downregulation of peptidoglycan synthesis, likely by inhibiting PstP’s dephosphorylation of peptidoglycan regulators (Figure 2,3,4). The mis-regulation of peptidoglycan metabolism (Figure 6c,d,e,f,g and Supplementary Figure S2b,d) may account for the slower growth of the *pstP*T134E*_Msmeg_* strains that we see in our data (Figure 6a), though it is likely that other processes beyond peptidoglycan metabolism are also misregulated, as PstP likely has many substrates involved in diverse processes (Molle *et al*., 2006; Iswahyudi *et al*., 2019).

### PstP localizes at the septum in exponentially growing *Msmeg* cells

We sought to determine if PstP’s activity against its diverse substrates could be regulated through protein localization. We made an *Msmeg* strain carrying an additional copy of *pstPWT_Msmeg_* fused to *mCherry2B*. By time-lapse fluorescence microscopy, we observed that mCherry2B-PstPWT*_Msmeg_* appears to be uniformly distributed along the membrane throughout most of the cell cycle, but becomes localized at the septum before cell separation (Figure 7). These data suggest that PstP is also regulated by its access to substrates at different sites in the cell.

**Figure 7.**
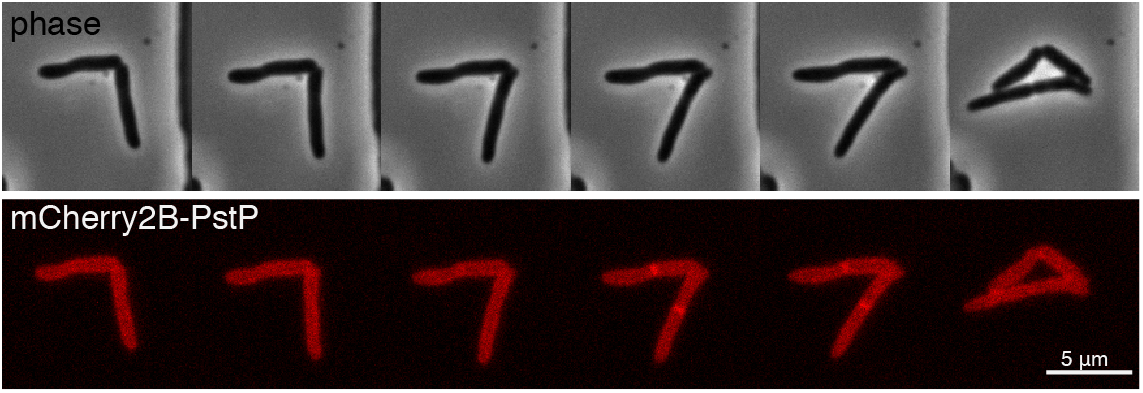
PstPWT*_Msmeg_* localizes at mid-cell. *Msmeg* cells expressing mCherry2B-PstPWT*_Msmeg_* were imaged over time in a CellAsic Microfluidic device at 15-minute intervals. Intensity is seen at the mid-cell (white arrows) for several frames before cell separation can be visualized by phase contrast microscopy (yellow arrows).

## Discussion

Our findings indicate that PstP has different basal activity rates against different substrates (Figure 4c), is regulated by its own phosphorylation (Figure 2,3,4,5) to regulate cell wall metabolism (Figure 6), and that it may also be regulated by its localization (Figure 7).

We see that PstPWT*_Mtb_* dephosphorylates CwlM*_Mtb_*, FhaA*_Mtb_* and Wag31*_Mtb_* at different rates *in vitro* (Figure 4c), with Wag31*_Mtb_* being the fastest to be dephosphorylated and FhaA the slowest (Figure 4c). We showed that PstP localizes at the septum, or cell division site, in the exponentially growing *Msmeg* cells (Figure 7). Both FhaA and Wag31 localize at the poles and septum (Gee *et al*., 2012; Hannebelle *et al*., 2020). Our localization data (Figure 7) suggest that co-localization of PstP with these substrates at the septum may cause both Wag31 and FhaA (Figure 2, 3, 4c) to be more highly phosphorylated at the cell poles than at the septum. As PstP dephosphorylates Wag31 so rapidly (Figure 3, 4c), we hypothesize that Wag31 is likely to be largely unphosphorylated at the septum, and may only be phosphorylated at the poles.

Our data in Figure 2, 3 and 4 show that negative charges at PstPT141*_Mtb_* and T174*_Mtb_* slow dephosphorylation of Wag31*_Mtb_*, but not FhaA*_Mtb_* and CwlM*_Mtb_*. Phosphorylation site T141 is on the flap subdomain and T174 is on the β-sheet core (β8) on PstP*_Mtb_* (Pullen *et al*., 2004), but both are facing into the same groove between these domains (Figure 1). T137 is the first residue of the flap subdomain (Pullen *et al*., 2004), and it faces away from the groove with the other phospho-sites (Figure 1). We propose that the T141/ T174 groove may contribute to toggling specificity of PstP against certain substrates, depending on the phosphorylation status.

A previous study showed that PstP_c*Mtb*_, phosphorylated by co-expression with PknA*_Mtb_* or PknB*_Mtb_*, showed higher activity than unphosphorylated PstP against a non-biological small molecule substrate *in vitro*, while PstP_c_T141E*_Mtb_* showed the same activity as the wild-type against the small molecule (Sajid *et al*., 2011). However, the status of phosphorylation of the other individual phospho-sites on PstP*_Mtb_* was not examined in that study. Our study, in contrast, sheds light on the effect of individual phospho-sites against biological protein substrates. Our findings (Figure 2,3, and 4) show that the phospho-mimetic PstP_c_T137E*_Mtb_* has an inhibitory effect whereas the phospho-mimetic variants PstP_c_T141E*_Mtb_* and PstP_c_T174E*_Mtb_* play a role in substrate specificity.

Our *in vitro* data in Figure 2, 3, 4 and 5 show that PstPT137E*_Mtb_* inhibits its activity against CwlM*_Mtb_*, FhaA*_Mtb_* and Wag31*_Mtb_*, and also against the STPKs PknA and PknB that phosphorylate them, all of which are involved in regulation of peptidoglycan metabolism (Thakur and Chakraborti, 2006; Kang *et al*., 2008; Jani *et al*., 2010; Gee *et al*., 2012; Prisic and Husson, 2014; Boutte *et al*., 2016; Viswanathan *et al*., 2016; Viswanathan *et al*., 2017; Carette *et al*., 2018; Turapov *et al*., 2018). The corresponding residue in PstP*_Msmeg_* is T134. Although this residue may not be phosphorylated in PstP*_Msmeg_* (Iswahyudi *et al*., 2019; Le *et al*., 2020), we sought to determine if a negative charge at this site would still affect peptidoglycan metabolism in *Msmeg*, as we would expect from the changes in dephosphorylation of the peptidoglycan regulators from *Mtb.* Our *in vivo* data show that the *pstP*T134E*_Msmeg_* strains grow more slowly, and have an increase in cell length and HADA staining, especially in starvation (Figure 6). This suggests that dephosphorylation of CwlM, FhaA, Wag31, PknB or PknA, is important in the transition to stasis and that a negative charge at the T134*_Msmeg_*/T137*_Mtb_* site would inhibit the ability of PstP to properly regulate the cell wall through these and other substrates. In this study, we focus on the residues that are phosphorylated in the PstP protein from *Mtb* (Sajid *et al*., 2011); while the experiments in *Msmeg* do not predict what effects phosphorylation on PstPT137 would have in *Mtb*, they at least indicate that some effect on peptidoglycan metabolism is probable.

Other studies have shown that PstP is an important regulator of cell division and cell length (Sharma *et al*., 2016). Our work shows that PstP localizes to the cell division site, which corroborates its importance in septation (Figure 7). We also show that the PstPT134E*_Msmeg_* mutation affects cell length (Figure 6b), which could be mediated by changes in division or elongation.

Although our findings are limited to PstP’s activity on certain substrates only, they show that the individual phosphosites are important in regulating PstP. *In vitro*, we see that a negative charge on T137 of PstP*_Mtb_* inhibits its activity against three PG regulators (Figure 2,3,4), as well as the essential kinases PknB and PknA when they are mixed together (Figure 5). In the cell, the substrates and kinases also co-exist, and PstP may exert its effect directly by dephosphorylating the substrates or indirectly by acting on the kinase (Bhaskara *et al*., 2019), or both simultaneously. There are likely other mechanisms of cell wall regulation involved in the complex *in vivo* scenario like protein localization or PstP’s interaction with other regulatory proteins that may contribute to PstP’s regulation as well. We do not know the conditions under which PstP is phosphorylated, or the combinations of PstP’s phosphorylations that exist in the cell. So the *in vitro* data only gives us a hint of the deep complexity of the STPK-substrate-PstP network *in vivo*.

In this work we show that PstP is regulated by both phosphorylation and localization. This work demonstrates the importance of protein dephosphorylation, not just phosphorylation, in regulating the activity of proteins involved in cell wall metabolism in mycobacteria. Future work that addresses when all of these important phospho-sites are phosphorylated and dephosphorylated will help us to understand how all these protein regulators contribute to coordinating the construction and integrity of the cell wall in changing conditions.

## MATERIALS AND METHODS

### Bacterial strains and culture conditions

All *Mycobacterium smegmatis* mc^2^155 ATCC 700084 cultures were grown in 7H9 (Becton, Dickinson, Franklin Lakes, NJ) medium containing 0.003 g/liter catalase, 5 g/liter bovine serum albumin (BSA), 2 g/liter dextrose, 0.2% glycerol, 0.85 g/liter NaCl, and 0.05% Tween 80 and incubated at 37°C until log. phase. 1xPBS supplemented with 0.05% Tween 80 was used for starvation assays. Cultures were plated on LB Lennox agar (Fisher BP1427-2).

*E. coli* Top10, XL1-Blue, and Dh5α strains were used for cloning and *E. coli* BL21 Codon Plus and BL21(DE3) strains were used for expressing proteins. Antibiotic concentrations for *M. smegmatis* were 25 μg/ml kanamycin and 20 μg/ml zeocin. Antibiotic concentrations for *E. coli* were 50 μg/ml kanamycin, 25 μg/ml zeocin, 20 μg/ml chloramphenicol, and 140 μg/ml ampicillin.

### Cloning and Strain construction

N-terminally Histidine (His) tagged cytosolic domains (1-300 amino acids (Gupta *et al*., 2009)) of *pstP*_c_T137E*_Mtb_* and *pstP*_c_T141E*_Mtb_*, and *fhaA*WT*_Mtb_* codon-optimized for expression in *E. coli* were ordered from IDT, Inc. and cloned in to pET28a.

The *Msmeg* mc^2^155 strains Δ*pstP*::lox L5::pCT94-p766tetON6-*pstP*T134E*_Msmeg_* and Δ*pstP*::lox L5::pCT94-p766tetON6-*pstP*WT*_Msmeg_* were generated by L5 allele swapping as described previously (Pashley and Parish, 2003; Shamma *et al*., 2021). All vectors and strains are described in supplementary tables 1-3.

A vector that expressed mCherry-2B-PstPWT*_Msmeg_* was constructed using isothermal assembly and integrated at the tweety phage integration site in *Msmeg*. In brief, *pstP*WT was amplified from *Msmeg* genomic DNA, and subcloned into a plasmid along with *mCherry2B* and the uv15 ribosomal promoter. The resulting vector was transformed into *Msmeg*.

### Growth Curve Assay

At least three biological replicates of *pstP*T134E*_Msmeg_* and WT allelic variants were grown in 7H9 medium up to the mid-log. phase. The growth curve assays were performed in non-treated non-tissue culture plates using a plate reader (BioTek Synergy neo2 multi mode reader) in 200μl of 7H9 medium at 37°C starting at an Optical Density at 600nm (OD_600_) of 0.1.

The doubling times of each strain were calculated in the non-linear (curve fit) method using an exponential growth equation with the least squares (ordinary) fit in GraphPad Prism (version 7.0d). *P* values were calculated using two-tailed unpaired t-tests.

### Protein purification

His-MBP-PknA*_Mtb_* was expressed in *E. coli* BL21(DE3) and all other proteins were expressed using *E. coli* BL21 Codon Plus cells. The *Mtb* genes of all studied factors were cloned into expression vectors. His-MBP-PknB*_Mtb_*, His-SUMO-CwlM*_Mtb_*, His_-_PstP_c_WT*_Mtb_* and His-PstP_c_T174E*_Mtb_* were expressed and purified as previously described (Kieser *et al*., 2015; Shamma *et al*., 2021). His-PstP_c_T137E*_Mtb_* and His-PstP_c_T141E*_Mtb_* were expressed and purified using the same conditions and buffers used to purify His-PstP_c_WT*_Mtb_* and His-PstP_c_T174E*_Mtb_* (Shamma *et al*., 2021). We purified His-PstP_c_WT*_Mtb_* and its phospho-mimetic variants using nickel affinity chromatography followed by Size Exclusion chromatography to ensure elution of the correctly folded proteins in the eluate. His-PstP_c_WT*_Mtb_* and all its phospho-mimetic variants are 34.1KDa and all of them eluted at the same column volume when run on the size exclusion resin (GE Healthcare Sephacryl S-200 in HiPrep 26/70 column). The purification chromatograms were analysed with a standard curve made from running protein standards (BIORAD) on the same size exclusion resin. All the PstP_c_ variants came out at a column volume where proteins weighing ∼35KDa elute according to our standard curve. This indicates that all the PstP variants fold like the wild-type.

His-Wag31*_Mtb_* was induced with 0.5mM IPTG (isopropyl-*β*-D-thiogalactopyranoside) for 6h at 18°C, and purified on Ni-NTA resin (G-Biosciences 786-940) then dialyzed and concentrated. The buffer for His-Wag31*_Mtb_* was 50mM Tris pH 8, 350mM NaCl, 1 mM dithiothreitol (DTT), and 10% glycerol. Imidazole (10mM and 20mM) was added to the buffers for lysis of the cell pellets and washing the Ni-NTA resin respectively, and 250mM imidazole was added for elution. The lysis buffer was supplemented with 1mM PMSF and 0.2% Triton X100.

His-MBP-PknA*_Mtb_* (1-279 amino acids containing the kinase domain (Baer *et al*., 2014)) was induced with 0.5mM IPTG at 18°C overnight, purified with 5ml Ni-NTA resins (BIO-RAD EconoFit Nuvia IMAC, 12009287), then run over the size exclusion resin (Enrich SEC650, BIO-RAD 780-1650) after dialysis and concentration to get soluble protein. The buffer for purifying His-MBP-PknA*_Mtb_* was 50mM Tris pH 7.5, 150mM NaCl, 20% glycerol. PMSF (1mM) was added to the buffer while resuspending the cell pellets. Imidazole was used in buffer for application to (20mM) and for elution from Ni-NTA resin (200mM).

His-FhaA*_Mtb_* was induced with 1mM IPTG at 18°C overnight and purified on Ni-NTA resin (G-Biosciences, 786-940 in 5 ml Bio-Scale Mini Cartridges, BIO-RAD 7324661), then dialyzed, concentrated, and run over size exclusion resin (GE Healthcare Sephacryl S-200 in HiPrep 26/70 column). His-FhaA*_Mtb_* was purified using the buffers described in (Roumestand *et al*., 2011).

### *In vitro* dephosphorylation assays

His-SUMO-CwlM*_Mtb_* at 0.15 μg/μl (2.64 μM) was phosphorylated *in vitro* with His-MBP-PknB*_Mtb_* at 0.015 μg/μl for 30 minutes at room temperature with 0.15μCi/μl ATP [γ-^32^P] (5×10^-8^ μmol) (3000Ci/mmol, 10mCi/ml, PerkinElmer BLU002A250UC) and 2mM MnCl_2_ in buffer (50mM Tris, pH 7.5, 250mM NaCl, 10% glycerol and 1mM DTT). Right after collecting kinase reaction samples, the leftover mixture was split into equal parts in different microcentrifuge tubes to carry on phosphatase assays by adding replicates of His-PstP_c_WT*_Mtb_*, His-PstP_c_T137E*_Mtb_*, His-PstPcT141E*_Mtb_* or His-PstPcT174E*_Mtb_* at 0.012 μg/μl to the kinase reaction mixtures containing 0.12 μg/μl of His-SUMO-CwlM*_Mtb_* and incubated in phosphatase buffer (50mM Tris pH 7.5, 10mM MnCl_2,_ and 1mM DTT) at room temperature up to 40 minutes. His-FhaA*_Mtb_* at 0.15 μg/μl (2.54 μM) was phosphorylated *in vitro* with His-MBP-PknB*_Mtb_* at 0.015 μg/μl for 45 minutes at 37°C with 0.15μCi/μl ATP [γ-^32^P] in presence of 5mM MnCl_2_ in 25mM Tris pH 7.5 and 1mM DTT. Right after collecting 0 minute kinase reaction sample, the reaction mixture was then split into equal parts and incubated with either PstP_c_WT*_Mtb_* or different phospho-mimetic variants of it at 0.012 μg/μl up to 80 minutes at room temperature in presence of the phosphatase buffer where the concentration of the substrate was 0.12 μg/μl For CwlM and FhaA, the amount of kinase and phosphatases was 1/10^th^ the amount of substrate, by weight.

His-Wag31*_Mtb_* at 0.15 μg/μl (4.90 μM) was phosphorylated with His-MBP-PknA*_Mtb_* at 0.03 μg/μl for 2 hours at room temperature in presence of 0.2μCi/μl ATP [γ-^32^P] (6.67×10^-8^ μM), 10mM MnCl_2_ and buffer (50mM Tris pH 7.5, 20% glycerol and 150mM NaCl). The amount of PknA was 1/5^th^ the amount of Wag31 substrate, by weight.

The kinase reaction mixture was equally split into separate tubes after the 0 second kinase reaction sample was taken. His-PstP_c_WT*_Mtb_* or its different phospho-mimetic variants at 0.0013 μg/μl were added to the mixture containing 0.13 μg/μl substrate and incubated at room temperature for up to 120 seconds in the phosphatase buffer. The amount of phosphatase was 1/100^th^ the amount of Wag31 substrate in the phosphatase reactions, by weight.

Control samples with no kinase and with no phosphatase were performed in all the above-mentioned substrate dephosphorylation assays. All *in vitro* dephosphorylation assays were carried out separately with two individually purified batches of WT and phospho-mimetic mutant of PstP as biological replicates. In all assays, the substrates were in molar excess over the ATP, so that all the ATP will have been consumed before the phosphatase is added, and re-phosphorylation by the kinases should not occur.

### Quantification of γ-^32^P signals

Samples from each *in vitro* biochemical reaction were collected before (0 minute or second) and after addition of each of the phosphatases at several different time points. The samples were run on 12% (Mini-Protean TGX, BIO-RAD, 4561046) gels and stained with Coomassie blue solution. After destaining, gels were dried using the GD200 Vacuum Gel Drying system (Hoefer Inc.) and imaged with phosphor screens (Molecular Dynamics) and a scanner (Storm 860, Amersham Biosciences). ImageQuant software (Molecular Dynamics) was used to process the images.

The protein bands on the gel and the phospho-signals on the autoradiogram were then quantified with FIJI. The intensities of the substrate protein bands on the Coomassie-stained gel and their corresponding phospho-signals on the autoradiograms at each time point were normalized against the respective protein band and the phospho-signal at 0 minute (His-SUMO-CwlM*_Mtb_* and His-FhaA*_Mtb_*) or 0 second (His-Wag31*_Mtb_*) before addition of the phosphatase. These relative intensities were used to calculate the intensity of phospho-signal over the amount of substrate for each time point. These values were used to plot the graph. The γ-^32^P signal intensities of the kinases on autoradiograms at each time point were normalized against the respective phospho-signal on the kinase at 0 minute (His-MBP-PknB*_Mtb_*) or 0 second (His-MBP-PknA*_Mtb_*) before addition of the phosphatase. These relative γ-^32^P signal intensity values were then used to plot the graph in GraphPad Prism (version 7.0d). All values were plotted and p-values were calculated using two-tailed, unpaired Student’s t-test in GraphPad Prism (version 7.0d).

### Cell staining

At least three biological replicates of *pstP*T134E*_Msmeg_* and WT*_Msmeg_* allelic variants were grown in 7H9 medium up to mid log. phase. Cultures were washed and then starved in 1xPBS with 0.05% Tween 80 (PBST) at OD_600=_ 0.3 for 15 minutes. 100μl of starved culture was then treated with 3μl of 10mM fluorescent D-amino acid HADA for 10 minutes. Cells were then pelleted and resuspended in PBST and fixed with 10μl of 16% paraformaldehyde for 10 minutes at room temperature. Cells were then pelleted and resuspended in PBST before imaging on agarose pads.

For staining cells in the log. phase, 1μl of 1mM HADA was added to 100μl culture grown up to the mid-log. phase and incubated at 37°C for 15 minutes. Stained cells were then pelleted, washed and fixed as described above.

### Static microscopy and image analysis

A Nikon Ti-2 widefield epifluorescence microscope with a Photometrics Prime 95B camera and a Plan Apo 100x, 1.45 NA objective lens was used to image cells. A 350/50nm excitation filter and a 460/50nm emission filter were used to take the blue fluorescence images for HADA staining. All images were captured using the NIS Elements software and analyzed using FIJI and MicrobeJ (Ducret *et al*., 2016). Appropriate parameters for length, width and area were set in MicrobeJ to detect cells. All detected dividing (V-snapping) cells in the images were split at the septum so that each daughter cell could be considered as a single cell. No overlapping cells in the images were included in the analysis.

Length and mean-intensities of HADA signals of at least 330 cells from each of *pstP*T134E*_Msmeg_* and *pstP*WT*_Msmeg_* (at least 110 cells from each of three biological replicate strains of each genotype) were quantified using MicrobeJ. The values of the mean intensities of 330 cells of each *pstP* allelic mutant and WT are represented in the graphs generated using GraphPad Prism (version 7.0d).

The medial intensity profiles of HADA signals in cells from different *pstP* allele strains in log. phase and starvation analyzed with MicrobeJ were plotted on the Y-axis over relative positions of cells using the “XStatProfile” plotting feature in MicrobeJ to show the subcellular localization of fluorescent intensities in cells. Demographs of HADA signal intensity across cell lengths in log. phase and starvation were built using the “Demograph” feature of MicrobeJ by plotting the medial intensity profiles of HADA signals.

### Time-lapse microscopy

An inverted Nikon Ti-E microscope was used for the time-lapse imaging. An environmental chamber was maintained at 37°C. Cells were cultivated in an B04 microfluidics plate from CellAsic; continuously supplied with fresh medium; and imaged every 15 minutes using a 60x 1.4 N.A. phase contrast objective, with an intermediate magnification of 1.5x. mCherry2B was excited with a SpectraX Illuminator at 561-nm. Fluorescence was detected with a bandpass filter of 595/40 and detected with an ORCA Flash Digital sCMOS.

#### Western Blotting

To verify expression of PstPWT*_Msmeg_*, T134A*_Msmeg_* and T134E*_Msmeg_*, cultures were grown in 10ml of 7H9 medium to OD_600_ = 0.8, pelleted and resuspended in 500μl PBS with 1mM phenylmethylsulfonyl fluoride (PMSF), and lysed (MiniBeadBeater-16, model 607, Biospec). The supernatants of the cell lysates were run on 12% resolving Tris-glycine gels and then transferred onto polyvinylidene difluoride (PVDF) membranes (GE Healthcare). To detect PstP-strep-tag II, rabbit anti-Strep-tag II antibody (1:1,000, Abcam, ab76949) in Tris-buffered saline supplemented with Tween 20 (TBST) and 0.5% milk and goat anti-rabbit IgG (H1L) horseradish peroxidase (HRP)-conjugated secondary antibody (1:10,000, Thermo Fisher Scientific 31460) in TBST were used.

## Acknowledgement

This work was supported by NIH grant R01AI148917 to CCB and R01AI148255 to EHR.

## Notes

### Competing Interest Statement

The authors have declared no competing interest.

