## Supplemental files for "Mycobacterial Serine/Threonine phosphatase PstP is phospho-regulated and localized to mediate control of cell wall metabolism"

### Supplemental Figures:

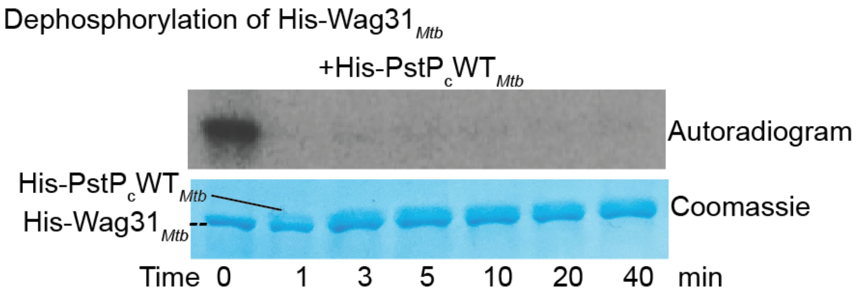

**Supplementary Figure S1: Dephosphorylation of His-Wag31*Mtb* by His-PstPcWT*Mtb*.** Autoradiogram and the respective Coomassie stained SDS-gel of *in vitro* phosphatase assays performed with ATP-[γ-$^{32}$P]-phosphorylated His-Wag31*Mtb* and His-PstPcWT*Mtb*. **His-PstPcWT*Mtb* used was 1/10$^{th}$ of** His-Wag31*Mtb* by weight in tthis assay.

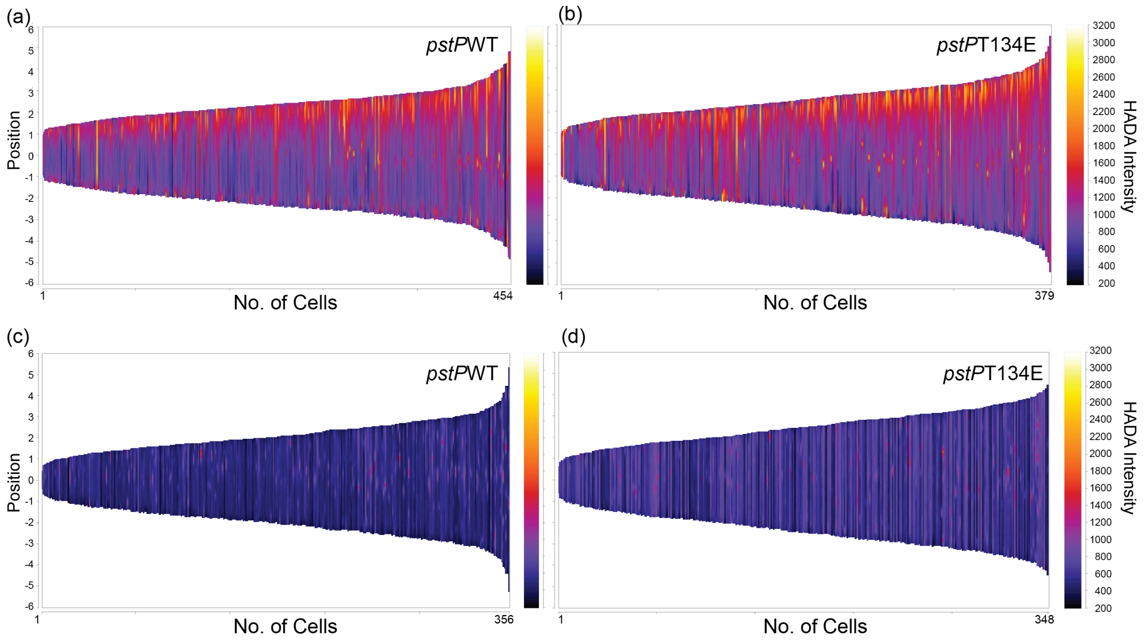

**Supplementary Figure S2: Demographic profiles of HADA signal intensity in *pstP*Msmeg strains in growth and starvation.** (a) and (b) Demographs showing intensities of the fluorescent dye HADA signal in individual cells from *pstP*WT*Msmeg* allele strains (a) and *pstP*T134E*Msmeg* allele strains (b) in log. phase.

(c) and (d) Demographs showing intensities of the fluorescent dye HADA signal in individual cells from *pstP*WT$_{Msmeg}$ allele strains (c) and *pstP*T134E$_{Msmeg}$ allele strains (d) starved in PBS with Tween80 for a total time period of 30 minutes. The left y-axis represents the position of septa and the poles of the cells. Cells are aligned at the septa (denoted by 0) with the fast growing old (brightest) old poles positioned above the septa and the slow growing new (dimmer) poles positioned below the septa. Different points in the cells are represented by positive units starting from the septa to the brightest (old) poles (0 to 6) and by negative units from the septa to the dimmer (new) poles (0 to -6). The right y-axis represents the intensity values of the HADA signal.

Biological triplicates of each *pstP*$_{Msmeg}$ allelic variant were analyzed. Signal intensities from at least 340 cells from each *pstP* allelic variant (at least 100 cells from each biological triplicate of each genotype) in log. phase and starvation were plotted in the demographs.

Dephosphorylation of His-SUMO-CwlM$_{Mtb}$

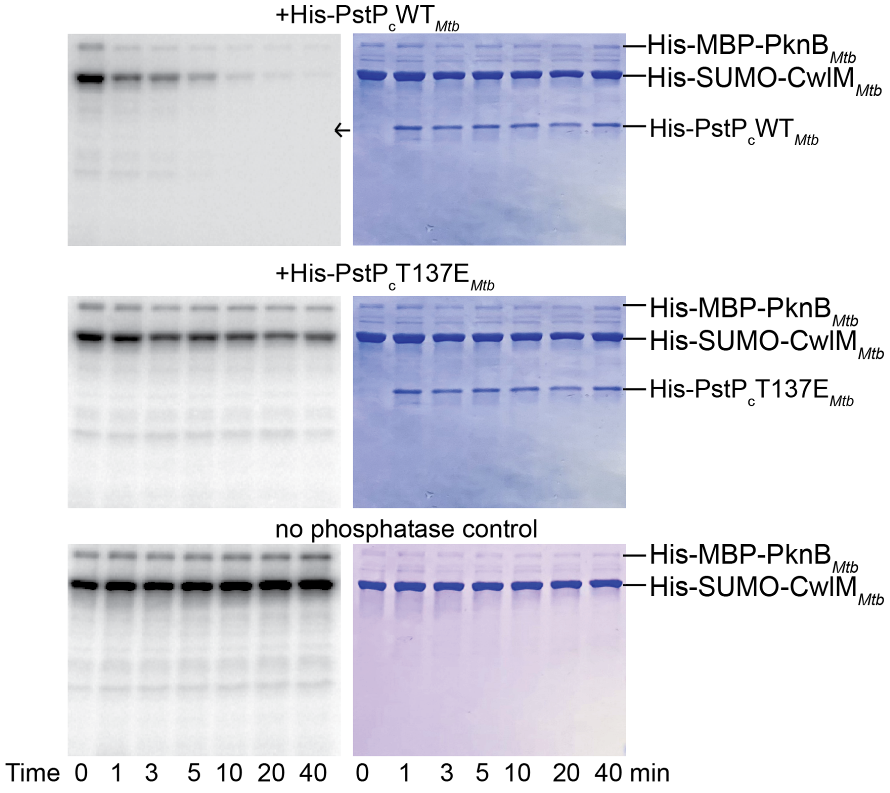

Dephosphorylation of His-Wag31$_{Mtb}$

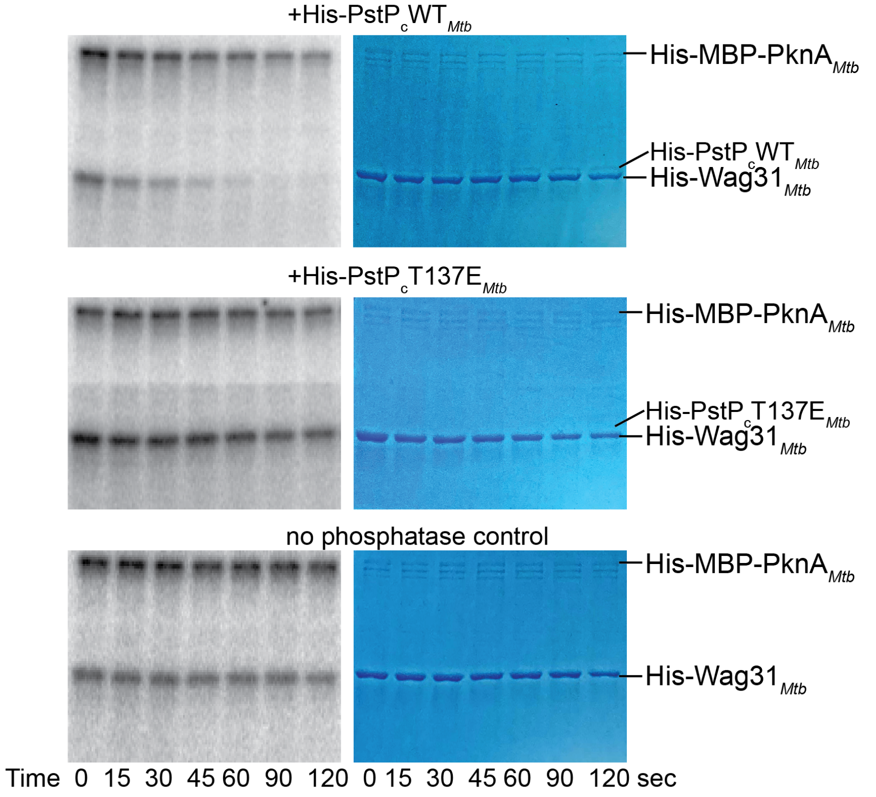

**Supplementary Figure S3: Dephosphorylation of His-SUMO-CwlM$_{Mtb}$ and His-Wag31$_{Mtb}$**

Top: Coomassie stained gels and the respective autoradiograms showing [γ-$^{32}$P]-phospho-signals on His-MBP-PknB$_{Mtb}$ and His-SUMO-CwlM$_{Mtb}$ in the *in vitro* phosphatase assays with His-SUMO-CwlM$_{Mtb}$ and His-PstP$_c$WT$_{Mtb}$ (first panel), His-PstP$_c$T137E$_{Mtb}$ (second panel) and no phosphatase (bottom panel).

Bottom: Coomassie stained gels and the respective autoradiograms showing [γ-$^{32}$P]-phospho-signals on His-MBP-PknA$_{Mtb}$ and His-Wag31$_{Mtb}$ in the *in vitro* phosphatase assays with His-Wag31$_{Mtb}$ and His-PstP$_c$WT$_{Mtb}$ (first panel), His-PstP$_c$T137E$_{Mtb}$ (second panel) and no phosphatase (bottom panel). One set of representative images for each reaction is shown here from two individual assays performed with individually purified batches of phosphatases.

**Supplemental Table 1: Strain list**

| Strain No. | Nick name | Genotype | Figure Panel |
|---|---|---|---|
| CB2378-79<br><br>CB2383 | *pstP*$_{Msmeg}$T134E slow growers | mc2155 ΔPstP::lox L5::pCT94-p766tetON6-PstPsmegT134E | 5a-g |
| CB2380-82 | *pstP*$_{Msmeg}$T134E | mc2155 ΔPstP::lox L5::pCT94-p766tetON6-PstPsmegT134E | 5a |
| CB1437-38, CB1292 | *pstP*$_{Msmeg}$WT | mc2155 ΔpstP::lox L5::pCT94-p766tetON6-PstPSmegWT | 5a-g |
| CB1755 | His-MBP-PknB$_{Mtb}$ | BL21CodonPlus/ pHMgWA-PknB(Kinase Domain) | 2a, 4a |
| CB1066 | His-MBP-PknA$_{Mtb}$ | BL21(DE3)/ pHMgWA-PknA (Kinase Domain) | 3a |
| CB1709 | His-SUMO-CwlM$_{Mtb}$ | BL21CodonPlus/pET-his-SUMO-CwlM | 4a |
| CB1710 | His-PstPcWT$_{Mtb}$ | BL21CodonPlus/pET28-his-PstPcWTMtb | 2a, 3a, 4a |
| CB1711 | His-PstPcT174E$_{Mtb}$ | BL21CodonPlus/pET28-his-PstPcT174EMtb | 2a, 3a, 4a |
| CB2498 | His-PstPcT137E$_{Mtb}$ | BL21CodonPlus/pCT216-his-PstPcT137Emtb | 2a, 3a, 4a |

| | | | |
|---|---|---|---|
| CB2499 | His-PstPcT141E$_{Mtb}$ | BL21CodonPlus/pCT216-his-PstPcT141Emtb | 2a, 3a, 4a |
| CB1287 | His-Wag31$_{Mtb}$ | BL21CodonPlus/ pET28 -WAG31(TB) | 3a |
| CB1962 | His-FhaA$_{Mtb}$ | BL21CodonPlus/pCT216-his-FhaAWTMtb | 2a |
| | m | | |

**Supplemental Table 2: Plasmid list**

| **Boutte Lab Strain # (Plasmid (p) in the strain)** | **Plasmid name** | **Used in strains** | **Reference for parent vector** |
|---|---|---|---|
| CB1354 | *pstP*$_{Msmeg}$T134E | CB2378-83 | (Shamma *et al.*, 2021) |
| CB1285 | p1210-p766TetON6-*pstP*$_{Msmeg}$WT | CB1437-38, CB1292 | (Shamma *et al.*, 2021) |
| CB174 | pL5 PTetO Msm PonA1 truncation A-FLAG clone 1 | all of the strains above | (Kieser *et al.*, 2015) |
| CB1155 (pCT298) | pET-his-SUMO-CwlM | CB1709 | (Shamma *et al.*, 2021) |
| CB1305 (pCT216) | pET28-his-PstPcWT$_{Mtb}$ | CB1710 | (Shamma *et al.*, 2021) |
| CB1532 (pCT216) | pET28-PstP(TB cyto) T174E | CB1711 | (Shamma *et al.*, 2021) |
| CB1069 | PHMgWA-PknB(KD) | CB1755 | (Boutte *et al.*, 2016) |
| CB1295 | His-MBP-PknA$_{Mtb}$ | CB1066 | (Baer *et al.*, 2014) |

| | | | |
|---|---|---|---|
| CB1281 | His-Wag31$_{Mtb}$ | CB1287 | This paper |
| CB1961 (pCT216) | His-FhaA$_{Mtb}$ | CB1962 | This paper |
| CB1786 (pCT216) | His-PstPcT137E$_{Mtb}$ | CB2498 | This paper |
| CB1787 (pCT216) | His-PstPcT141E$_{Mtb}$ | CB2499 | This paper |

**Supplemental Table 3: Primer list**

| Plasmid in Strain # | Feature | Primer | Primer # | Reference |
|---|---|---|---|---|
| CB1354 | pCB1285-p766tetON6-PstP$_{Msmeg}$T134E-strep | tttcttgtacaaagtggctctcctCATatgaccctcgttctccgctac | FS 40 | (Shamma *et al.*, 2021) |
| | | acagatcaccaaggacgacGAGttcgtgcagaccctcgt | FS 41 | |
| | | acgagggtctgcacgaaCTCgtcgtccttggtgatctgt | FS 42 | |
| | | TAGGGTCCCCAATTAATTAGCTAAAGCTTtcaCTTCTCGAACTGGGGGTGGCTCCAGTCG | FS 12 | |
| CB1285 | p1210-p766TetON6-PstP$_{Msmeg}$WT-strep | TGCTTAATTAAGAAGGAGATATACATatgaccctcgttctccgctacg | FS 10 | (Shamma *et al.*, 2021) |
| | | GAACTGGGGGTGGCTCCAGTCGGCGCCGGTGGAGTGtgacaccgcccggcagttcgtccc | FS 11 | |
| | | TAGGGTCCCCAATTAATTAGCTAAAGCTTtcaCTTCTCGAACTGGGGGTGGCTCCAGTCG | FS 15 | |
| | | CGTTTAATACTGCATGCACTCTAGAgctaccaggcctagatctggggacc | FS 21 | |
| | | cgtagcggagaacgagggtcatATGaggagagccactttgtacaagaaag | FS 22 | |
| | | tttcttgtacaaagtggctctcctCATatgaccctcgttctccgctac | FS 40 | |
| | | ggccacgaggtcgagccgGAGctgatcatgcgcgaggcc | FS 45 | |

| | | | | |
|---|---|---|---|---|
| | | ggcctcgcgcatgatcagCTCcggctcga cctcgtggcc | FS 46 | |
| CT216 | pET28-his-PstPcWT*Mtb* | cctggtgccgcgcggcagccatatggcgcg cgtgaccctggtcctgcgat | FS 33 | (Shamma *et al.*, 2021) |
| | | gtggtggtggtggtggtgctcgagtcaccgtc ggcccgaccaccgtggcc | FS 34 | |
| CT216 | pET28-PstPc(TB cyto)T174E | cctggtgccgcgcggcagccatatggcgcg cgtgaccctggtcctgcgat | FS 33 | (Shamma *et al.*, 2021) |
| | | gtggtggtggtggtggtgctcgagtcaccgtc ggcccgaccaccgtggcc | FS 34 | |
| | | gttgaccggccatgaggtcgaaccgGAGc tgaccatgcgagaagcccgcgccggtgat | CB 1460 | |
| | | atcaccggcgcgggcttctcgcatggtcag CTCcggttcgacctcatggccggtcaac | CB 1461 | |
| CT298 | pET-his-SUMO-CwlM*Mtb* | Tcacagagaacagattggtggatccatgcc gagtccgcgccgcgaa | CB 1272 | (Shamma *et al.*, 2021) |
| | | gtgctcgacaagcttattactcgagttaagaa ccgccgagtctacc | CB 1273 | |
| KP35-79 | ΔpstP-hyg L5::p46-*pstPMtbWT*-strep | CGGATCGGCAAGACGGTAATC GAGCTGCGCCCGTGAGCCCG CGCACGCGAGGAGCAGACGC TCTAGAACTAGTGGATCC | S1256-SMEG-PstP-P1b | (Shamma *et al.*, 2021) |
| | | GGCGGCGTGACGGTAACCGG CGACTGGGGTTGCGTCGTCAT TCCTTCCTCCTTTCTTACTTCT AGACTCGAGGTACCG | S1257-SMEG-PstP-P2b | |
| | | CCAACGGCACTTACCTTGACA GGGCGAAGGTGACAACAGCA GTAAAGGTTCCCATTGGCGCG CCGGTGCGGATCGGCAAGAC GGTAATCG | S1258-SMEG-PstP-P3b | |
| | | TCGACGATCAACAGGGCCACG GTGGTGATCAGCGCCGCGAAC CCCAGCAGCAGCAGTTCCGCA TTGCGCCGGTTCGGGAGCGG CGGCGTGACGGTAAC | S1259-SMEG-PstP-P4b | |
| | | AACTCGACGGCATGGGCAC | S1260-SMEG-PstP-del Int-For2 | |
| | | GCGAGATGCCCAGGAAGGAG | S1261-SMEG-PstP-del Int-Rev2 | |
| | | GCGCGGTGATGGTGAGAC | S1121-SMEG-PstP del P6-ExtFor | |
| | | GCAGGCTCGCGTAGGAATCAT CC | S486-Hyg-N-out | |
| | | GAACTGCTCGCCTTCACCTTC C | S538-Hyg-C-out | |

| | | | | |
|---|---|---|---|---|
| | | TTCAAGAACACCACCACGAGC | S1122-SMEG-PstP del P7 ExtRev | |
| CB1281 | pET-his-Wag31$_{Mtb}$ | cctggtgccgcgcggcagccatatgatgccgcttacacctgccgacgtcc | NH 22 | This paper |
| | | agtggtggtggtggtggtgctcgagctagttttgccccggttgaattga | NH 23 | |
| | pTw-mcherry2B-PstP | cgagccgccACTGGATCCGCTAGATCC | mcherry 2b-linker-R | |
| | | GGATCTAGCGGATCCAGTggcggctcgaccctcgttctccgctacgcgg | mcherry 2b-pstp-3 | |
| | | GGGTCCCCAATTAATTAGCTAAAGCTTtcatgacaccgcccggcagttcg | mcherry 2b-pstp-4 | |
| | | GGCTCTGGGAGTACCCGTG | KB 414 For | |
